# Rare long-range cortical connections enhance information processing

**DOI:** 10.1101/2021.02.08.430236

**Authors:** Gustavo Deco, Yonathan Sanz Perl, Peter Vuust, Enzo Tagliazucchi, Henry Kennedy, Morten L. Kringelbach

## Abstract

What are the key topological features of connectivity critically relevant for generating the dynamics underlying efficient cortical function? A candidate feature that has recently emerged is that the connectivity of the mammalian cortex follows an exponential distance rule, which includes a small proportion of long-range high-weight anatomical exceptions to this rule. Whole-brain modelling of large-scale human neuroimaging data in 1003 participants offers the unique opportunity to create two models with and without long-range exceptions and explicitly study their functional consequences. We found that rare long-range exceptions are crucial for significantly improving information processing. Furthermore, modelling in a simplified ring architecture shows that this improvement is greatly enhanced by the turbulent regime found in empirical neuroimaging data. Overall, the results provide strong empirical evidence for the immense functional benefits of long-range exceptions combined with turbulence for information processing.

## Introduction

In Nature, complex dynamics of physical systems emerge from the integration of the underlying dynamics parts into the whole. In general, global integrated dynamics are shaped by the structure of the coupling between the parts, which in most cases are simple short-range local interactions. In neuroscience, structural investigations show that the brain is a particularly interesting physical system where much of the wiring is local but with important exceptions in long range white-matter connections ^1-9^. In fact, this apparent complexity has been shown by recent research using massive tract-tracing studies to be surprisingly simple: the anatomical architecture of the mammalian cortex uses similar simple short-range wiring with an exponential drop off in strength over distance, commonly known as the exponential distance rule (EDR) ^10-12^. Crucially, however, these investigations have also demonstrated that on top of EDR, there are rare long-distance connections (EDR+LR) which have a non-random role in the specificity and non-homogeneity of the cortical architecture ^1,2^. It is highly likely that these weight-distance relations make brain architecture unique among known physical systems.

However, importantly the functional relevance of these long-range exceptions to brain dynamics remains unresolved. It is clearly impossible to selectively remove these long-range exceptions in animals experiments which could allow a disentanglement of their functional role. Previous research has therefore investigated *in silico* the structural consequences of manipulating the topology of brain anatomy ^13-15^, and there have even been some studies of their functional consequences ^16,17^. Frustratingly specific information processing capability offered by the long-range exceptions remains unresolved.

Here, whole-brain models constructed according to the structural Bauplan of the brain and incorporating sophisticated local coupling are fitted to the empirical data and shown to replicate the global brain dynamics ^18-20^. This explicitly allow us to change the underlying anatomical structural connectivity to be either local EDR wiring or both local and long-range (EDR+LR) in order to uncover the functional role of the rare long-range exceptions.

The whole-brain model contains two main ingredients, namely anatomy and dynamics, which are used to accurately fit and reproduce many aspects of empirical neuroimaging data, including the functional connectivity, functional connectivity dynamics, metastability and even clustered composition of functional connectivity. The source code for these models are freely available and there is even The Virtual Brain (https://www.thevirtualbrain.org/). As can be seen from this code base, for estimating the anatomical structural connectivity, excellent results can be obtained from diffusion MRI ^21^. For modelling local dynamics, initially complex models of neural dynamics going from spiking neuronal circuits ^22^ to dynamical mean field ^23^ were used. More recently a more efficient and yet more simple, yet very effective model has been shown to be a mesoscopic system using the so-called Stuart-Landau oscillators ^24^. As an example of the power of the whole-brain models, the Hopf models is able to describe up to 80% of the resting state functional connectivity. Still, there are some important features of brain dynamics such as non-stationarity and non-equilibrium which the current models are not yet able to capture. However, this remains an active area of research ^25^.

Oscillators have been used to model many physical systems, going from the simplest linear, harmonic oscillator to non-linear oscillators ^26^. Small perturbations to linear oscillators lead to changes in oscillation amplitudes, while perturbations to non-linear oscillators lead to self-regulating relaxation and a return to the same region in phase space. With an ordinary differential equation of a complex order parameter, the Stuart-Landau model of a single oscillator provides the simplest nonlinear extension of a linear oscillator that mathematically describes the onset of spontaneous oscillations (i.e. bifurcation from fixed-point dynamics towards a limit cycle). This model has been shown to be remarkably effective at modelling the mesoscopic dynamics of brain regions, which contain combinations of different noisy to oscillatory regimes.

In neuroscience, the whole-brain model using dMRI and Stuart-Landau oscillators is called the *Hopf whole-brain model* in honour of the German mathematician Eberhard Hopf who described the normal form of the Hopf bifurcation, which describes the behaviour of a Stuart-Landau non-linear oscillating system ^27^. In physics, similar large-scale system of spatially coupled oscillators has been shown to give rise to a wealth of spatiotemporal patterns, ranging from regular laminar waveforms to highly turbulent dynamics. Interestingly, similar to these findings in physics, the Hopf whole-brain model has recently been used to capture the regularities of empirical neuroimaging data exhibiting turbulent dynamics ^28^. Using a large, high-quality state-of-art dataset of 1003 HCP participants it was shown that human brain dynamics exhibit turbulence a power law similar to that shown by Kolmogorov in fluid dynamics. Equally, the results show turbulence in the empirical data as formalised by Kuramoto in his studies of coupled oscillators. Furthermore, building a whole-brain Hopf model with Stuart-Landau coupled oscillators, it was demonstrated that maximal turbulence and information processing is found at the optimal working point of the model.

In terms of the physiology, the important principles for turbulence in the brain, as already demonstrated ^29^, is not the kinetic movement of fluids but the neural activity in terms of local synchronisation. This is the key insight of Kuramoto who was able to demonstrate that oscillators with local synchronisation can lead to turbulence in a non-fluid context when coupled in the right way ^30^. Even more, given that the brain can be described by coupled oscillators, this closes the circle.

Here we investigated the functional consequences of having EDR or EDR+LR anatomy in the Hopf whole-brain model fitted to very large-scale empirical neuroimaging data from 1003 participants in the human connectome project (HCP). First, we used diffusion (dMRI) tractographic data to identify the relationship between connection weight (in tractography referred to as streamline density) and distance. As expected, we found that the empirical tractographic connectivity data in the human cortex fits an EDR. Second, we identified rare long-range exceptions corresponding to those long-range connections with stronger weights than expected from the EDR. Third, whole-brain modelling was used to investigate the functional consequences of the EDR and EDR+LR models. We found that the addition of the long-range exceptions leads to a significant improvement to fitting the empirical neuroimaging data and, more importantly, a very significant enhancement of information processing, which we measured in four complementary ways.

Our first approach was to measure information processing by characterising the correlations between long-distance pairs of regions in MNI space. We then used three more sophisticated approaches which use the concept of ‘brain vortex space’, defined as the local level of synchronisation and thus similar but not identical to rotational vortices found in fluid dynamics.

The discoveries of Kuramoto of linking oscillators and turbulence, motivated our use of the concept of brain vortex space, which is exactly defined as the local level of synchronisation. This is thus similar but not identical to the rotational vortices found in fluid dynamics. In this case, the brain vortex space is the level of local synchronisation capturing the level of rotationality, whereas in fluid dynamics, the vortices are essentially capturing the rotational kinetic energy. Thus, while not identical, the brain vortex space shares some similarities with the vortex space used in fluid dynamics. Independent of semantic analogy, the concept of local synchronisation is fundamental for capturing the information transmission across spacetime. Indeed, based on this central concept we define three different measures, namely information cascade, susceptibility and predictability (see Results and Methods).

Finally, we investigated the role of turbulence for this significant improvement in information processing. This can be studied explicitly in the simplest possible ring structures of coupled Stuart-Landau oscillators with three different types of coupling: nearest neighbour, EDR, and EDR+LR. In this simpler model, we were able to modulate the levels of turbulence by changing the global coupling. We found that a low proportion of long-range exceptions critically are important to significantly improve the transmission of information across the system, i.e. the information cascade in a turbulent regime.

Overall, we found that a brain architecture with rare long-range exceptions play a crucial role for enhancing the information processing. We showed the long-range exceptions are important for building up resting state networks and as such could add a mechanistic underpinning for the recent systematic evidence of importance of long-range connectivity for cognition ^31,32^. The results demonstrate the functional benefits in terms of improving the information cascade provided by the unique brain architecture with EDR and long-range exceptions exploiting the turbulent-like regime of brain dynamics.

## Results

We hypothesised that adding the rare long-range exceptions to EDR crucially enhances information processing (as measured by our four measures of long-range connectivity, information cascade, susceptibility and predictability) over and above having only EDR as a wiring principle. While it is not possible to empirically remove them to test their functional consequence, we were able to use whole-brain modelling fitted to large-scale human neuroimaging data to explore their functional relevance.

The overall strategy is described in **Figure 1**. First, we fitted an EDR to human diffusion MRI and extracted rare long-range exceptions on top of these (EDR+LR). **Figure 1A** shows cartoons of the two different anatomical models: EDR model (blue) and EDR+LR model (red). **Figure 1B** shows how we built two different whole-brain models based on the two different anatomical models to test the differential functional role of rare long-range exceptions. We fitted both Hopf whole-brain models to the functional MRI data from 1003 participants. At the corresponding optimal working point, the models were able to reproduce the empirical whole-brain dynamics that emerges from the local dynamics of each brain region (described using a Stuart-Landau oscillator) coupled through the two different underlying anatomical hypotheses. As shown in **Figure 1C**, this framework allows us to test the functional consequences between the two models by using two sensitive measures of long-range functional connectivity (FC) and information cascade (see below and Methods) to assess the functional advantages of the rare long-range exceptions.

**Figure 1.**
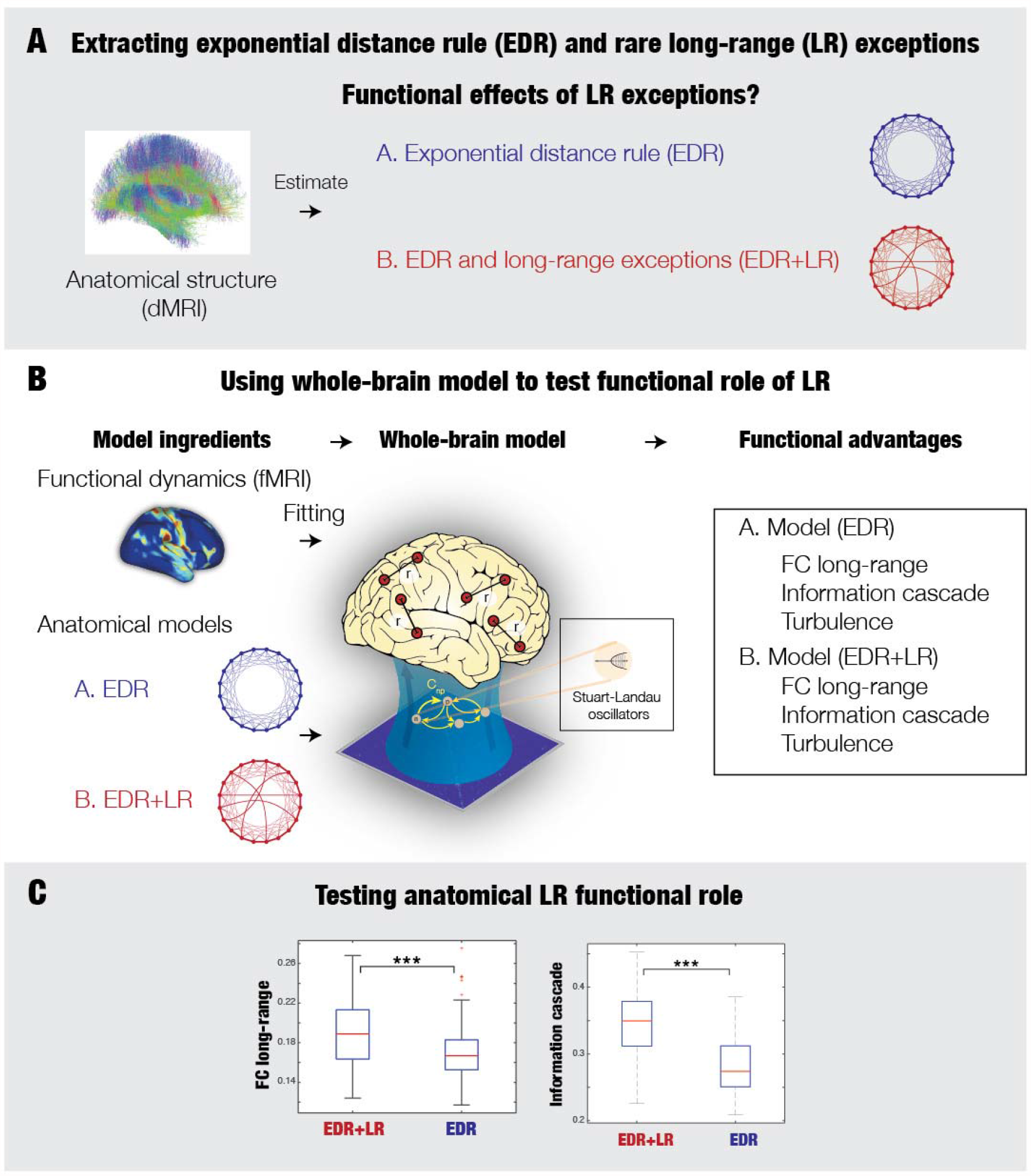
Using whole-brain modelling to determine the functional importance of recently discovered rare long-range exceptions. Massive tract-tracing studies in primates have revealed the simple, yet powerful economy of anatomy as a cost-of-wiring principle of rare long-range exceptions on top of an exponential distance rule. Whole-brain modelling provides a unique opportunity to disentangle the functional role of these long-range exceptions by testing alternative hypotheses. **A)** First, human diffusion MRI was used to fit the exponential distance rule (EDR) and extract rare long-range exceptions on top of these (EDR+LR). The two different anatomical hypotheses are illustrated by the cartoon of two rings (EDR in blue and EDR+LR in red). **B)** Second, we built two different whole-brain models using these anatomical hypotheses to fit the functional MRI data from 1003 participants. At the optimal working point, these Hopf models are able to reproduce the empirical whole-brain dynamics that emerges from the local dynamics of each brain region (described using a Stuart-Landau oscillator) coupled through the two different underlying anatomical hypotheses. **C)** We used the two sensitive measures of long-range FC and information cascade to assess the functional advantages of the rare long-range exceptions.

### Extracting two underlying anatomical models to be tested

The functional advantages of the rare long-range exceptions can be investigated with whole-brain modelling using either EDR or EDR+LR anatomical models. **Figure 2A** shows that the mammalian cortex is well-described by an architecture with long-range exceptions on top of an EDR ^11^. The anatomical structural connectivity of the human brain can be estimated using dMRI tractography. We estimated empirical HCP dMRI tractography of the human brain by estimating the streamline densities (i.e. connection weights) between the pairs of regions in the fine-grained Schaefer parcellations (with 1000 parcels) as a function of the Euclidean distance between nodes (**Figure 2B** and Methods). For each region pair, we computed the Euclidean distance, r, in MNI space (**Figure 2C**) in order to access the spatial information required to investigate the rules underlying coupled connectivity.

**Figure 2.**
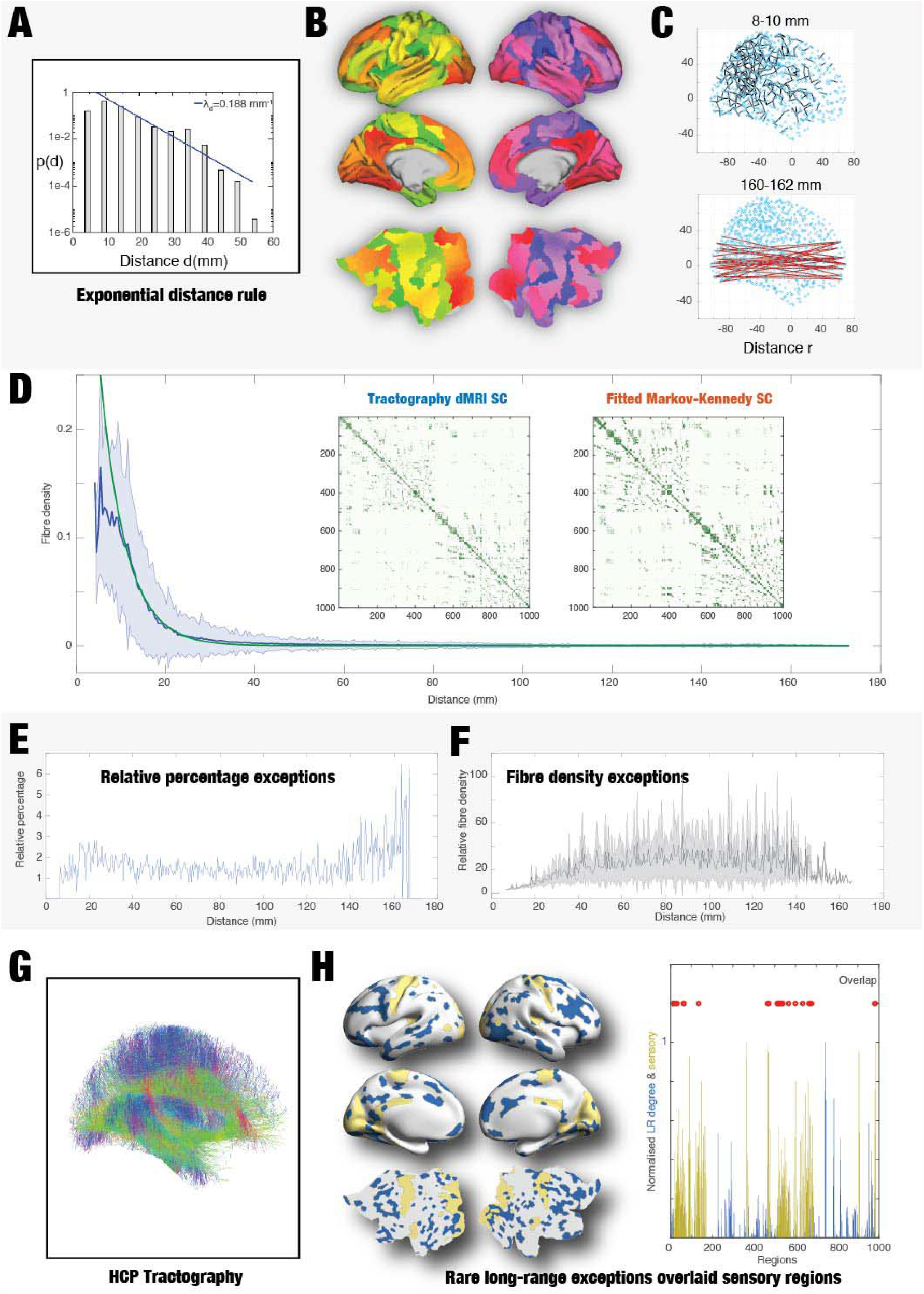
Extracting exponential distance rule (EDR) and rare long-range exceptions on top (EDR+LR). **A)** Consistent tract tracing studies in non-human primates have shown that most of the underlying brain connectivity follows the exponential decay described by the EDR ^11^. Here is shown the histogram of interareal projection lengths for all labelled neurons (n = 6,494,974), where the blue line shows the exponential fit with a decay rate 0.19 mm^-1^. **B)** In order to assess whether this EDR holds for the human brain, we used the fine-grained Schaefer parcellation with 1000 parcels, here shown as slices in MNI space and on the surface of the HCP CIFTI space. **C)** We computed the Euclidean distance, r, in MNI space between pairs of regions. Here we show two examples of the pairs with r=8-10 mm (top, black lines) and r=160-162 mm (bottom, red lines). **D)** We estimated the empirical HCP dMRI tractography of the human brain, as shown by the streamline densities between the pairs of regions in the Schaefer parcellations as a function of the Euclidian distance between the nodes. We found that the EDR is a good approximation of the human structural anatomical connectivity as shown by the red line showing the fitted EDR with an optimal λ=0.18 mm^-1^ fitting the empirical dMRI tractography (blue line). Equally, the remarkable similarity can be appreciated by comparing the two matrices showing the structural connectivity matrices for the empirical dMRI tractography (left subpanel) and the optimally fitted EDR connectivity (right subpanel). **E)** However, our results also demonstrate that similar to the non-primates, the anatomy is also characterized by a small proportion (1.23%) of long-range outliers of the EDR. We identify an exception by computing the distribution for a given distance and selecting those connections that are 3 standard deviations above the mean. Here, we plot the relative percentage of long-range outliers (for pairs at a given distance) as a function of that distance. Note the increase in relative percentage, especially for the longest-range connections. **F)** For these exceptions, we show relative streamline densities (for the pairs at a given distance) as a function of distance. Note the general tendency for an increase in the long-range connections. **G)** We plot a rendering of the combined HCP tractography in MNI space. **H)** The left panel shows a rendering on the human brain of the regions (in blue) as a degree of the matrix of the long-range EDR outliers overlaid with primary sensory regions (in yellow, as indexed by the myelin ratio). Right panel shows the normalised values for long-range EDR outliers (in blue) and sensory areas (in yellow) across regions, as well as red circles showing the 14.9% overlap between them.

**Figure 2D** shows the fitting of weights of connections as a function of distance, r, by an exponential decay function (with the shadow showing the standard deviation, see Methods). This means that the human structural anatomical connectivity exhibits a fitted EDR as reflected by the red line with an optimal λ_*S*_ = 0.18 mm^-1^ fitting the empirical mean dMRI connectivity matrix across participants (blue line). This is lower than the values λ_*S*_ = 0.78 mm^-1^ found in mice and similar to the value λ_*S*_ = 0.19 mm^-1^ found in non-human primates ^11,12^. Unlike the results in mice and non-human primates, the estimate found here is based on dMRI tractography and may be overestimating the λ_*S*_, which is likely to be smaller. In fact, fitting the functional neuroimaging data with λ_*S*_ as a free parameter yielded values lower than the one estimated here from the structural data^28^. Nevertheless, for consistency we use the latter here. The figure also shows the remarkable similarity between the two matrices representing the empirical dMRI structural connectivity matrix (left subpanel) and the optimally fitted EDR connectivity (right subpanel).

Although dMRI is not ideal for estimating precise anatomical connections ^33^, we designed an algorithm to identify rare long-range outliers to the EDR, i.e. connections that are much stronger than average as derived from the EDR. This algorithm first computes the distribution of weight connections at a given distance, r, in the average dMRI connectivity matrix. We then selected only those connection pairs that are three standard deviations above the mean weighting of connection at that given distance, r.

We found that similar to the cortical anatomy of non-human primates and rodents ^10-12^, human structural anatomy is characterized by a small proportion (1.23%) of rare long-range outliers of the EDR. To further specify the long-range exceptions over and above the standard deviation shown in **Figure 2D**, we are explicitly showing the relative percentage exceptions of long-range connections (for pairs at a given distance) as a function of that distance in **Figure 2E**. As can be clearly seen, long-range outliers increase in relative percentage with increasing distances.

Furthermore, **Figure 2F** shows the relative streamline density as a function of distance for the pairs of exceptions at a given distance. Note the general trend for an increase of relative streamline density in the long-range connections, again suggesting that the EDR outliers that are expected to play a predominant role in shaping dynamics are more prevalent at longer-distances.

For visualization, **Figure 2G** shows a rendering of the combined HCP tractography in MNI space (without cerebellum and brainstem). Crucially, the edge-complete connectome can be compared with the relative simplicity of the spatial location of the long-range outliers shown in **Figure 2H**. Here, we computed the regions with long-range exceptions (larger than 40mm, accounting for 78% of all) where the regions were computed as the degree of the long-range exception matrix. We render the pairs of regions (in blue) on various views of the human brain overlaid with the primary sensory regions (shown in yellow). The right panel shows the normalised values for each across the 1000 regions as well as red circles showing the small 14.9% overlap between them. This demonstrates that the long-range exceptions are mainly found in higher association cortex, which has been shown by many studies to be involved in higher brain function^34^.

### Rare long-range EDR outliers in human brain architecture improves information processing

We aimed to investigate the hypothesis the effect of rare long-range EDR outliers in information processing as measured by three measures of information transmission in brain vortex space and one measure of long-range correlation in MNI space. We used whole-brain models since they offer the advantage that they can be systematically altered to fit the empirical data, making it possible to test different hypotheses ^21^.

As shown in the previous section, from the empirical anatomical data we extracted two connectivity matrices, 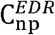, with the EDR and with long-range connections (larger than 40mm) on top (EDR+LR), 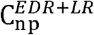 (see Methods). We used these for the anatomical connectivity matrices in two separate Hopf whole-brain models fitting empirical functional neuroimaging from 1003 human subjects (see Methods).

First, in order to show the appropriateness of our whole-brain model for hypothesis testing, we show the results of fitting to the empirical data. **Figure 3A** shows the evolution of the error of the FC fitting to the empirical data for both models as a function of the global coupling strength, G. The error of the FC fitting is given by the square root of the difference between the simulated and empirical FC matrices. We found for each model an optimal working point with G_EDR_=1.55 and G_EDR+LR_=1.3, both with a very good level of fit to the empirical data. We used these values as the basis of the following investigations to study our two models.

**Figure 3.**
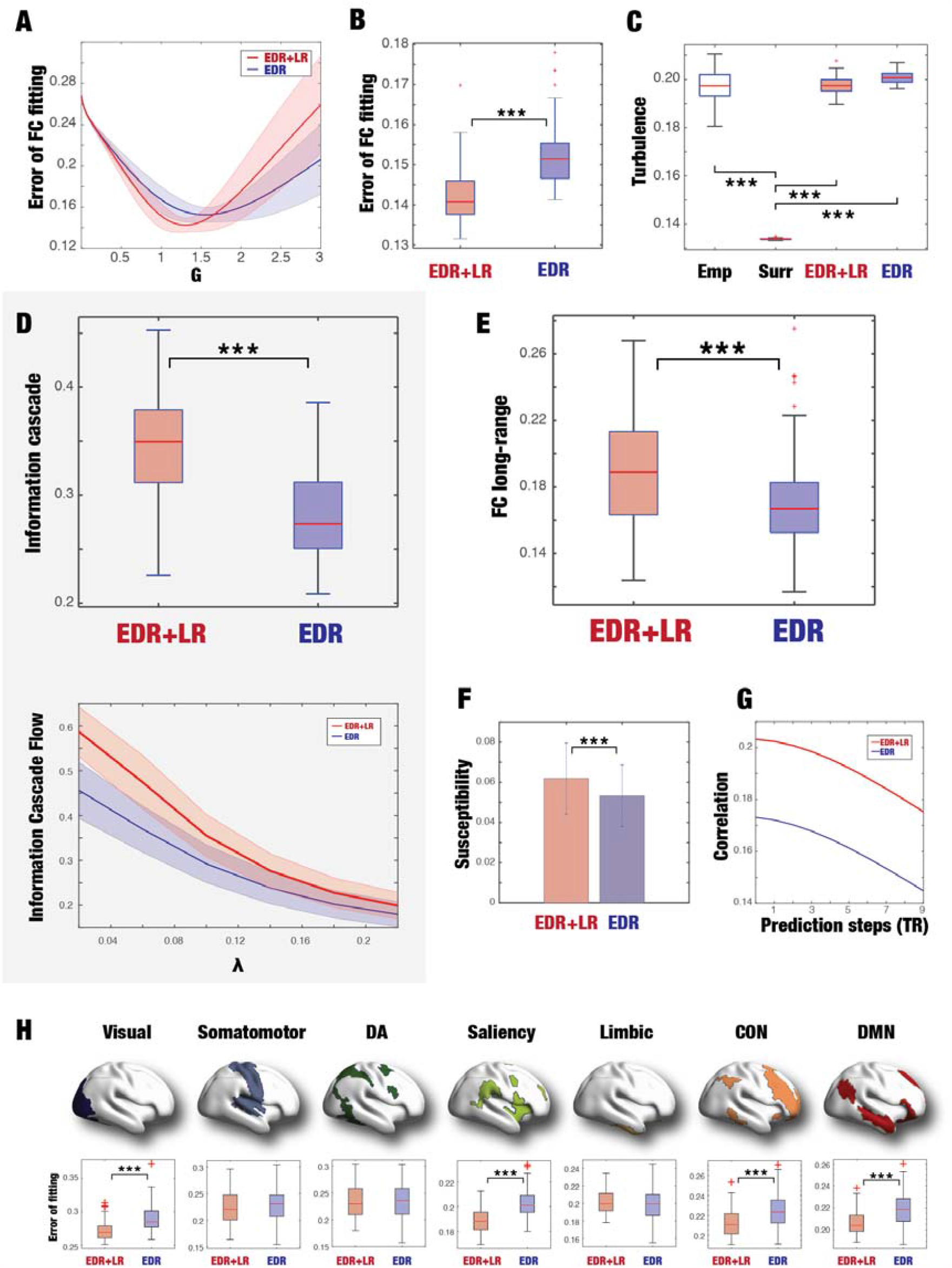
Whole-brain modelling shows how rare long-range exceptions in human brain improves information processing. As shown in Figure 2, we extracted two matrices with the exponential distance rule (EDR, blue) and EDR+long-range (EDR+LR, red) from the empirical human anatomical data. These two coupling matrices were used in a Hopf whole-brain model of Stuart-Landau oscillators fitting the empirical functional neuroimaging, and we chose the respective model optimum. We analysed the information processing ability of these two different architectures. **A)** The panel shows the error of the FC fitting to the empirical data for both models as a function of the global coupling strength, G. We use the respective minima (blue line, G_EDR_=1.55 and red line, G_EDR+LR_=1.3) as the basis of the following investigations. **B)** The panel shows boxplots of the errors of the FC fitting for the two models. The EDR+LR with the long-range connections perform significantly better (p<0.001, Wilcoxon rank sum). This shows the important role of the rare long-range exceptions. **C)** The panel shows that the amplitude turbulence for the EDR+LR and EDR models is both similar to the empirical (Emp) data – but not the surrogate (Surr) data. This suggests that the EDR+LR model is not affecting the simplest measure of turbulence data but as we show in Figure 4 (see text) turbulence does affect the role of LR exceptions in information processing. **D)** More specifically, we investigated the role of LR exceptions in information processing by measuring the information cascade (see Methods). The upper panel confirms the significant role (p<0.001, Wilcoxon rank sum) for LR exceptions in increasing the information cascade (compare the EDR+LR with EDR boxplots). The information cascade is the integration of information cascade flow across scales (see Methods). The bottom panel shows how the EDR+LR model is significantly higher than the EDR model in terms of information cascade flow as function of the scales in brain vortex space (compare red line for EDR+LR and blue line for EDR models). **E)** The panel shows boxplots of the mean values of the FC long-range (involving pairs with distances over 40mm) for the two models across 100 trials. There is a significant increase for the EDR+LR model (p<0.001, Wilcoxon rank sum), which shows the important role of long-range exceptions. **F)** Using whole-brain modelling allows to measure the susceptibility (the reaction of the model to external perturbation, see Methods) of the two models and again the EDR+LR model outperforms the EDR model. **G)** We also measured the predictability in brain vortex space for n steps in the future (shown on x-axis). Again, the EDR+LR model outperforms the EDR model. **I)** The seven subpanels show brain renderings of the seven Yeo resting state networks with the corresponding boxplots of the FC fitting of the two models (smaller values are better). Again, the EDR+LR model outperforms the EDR model and is able to better fit the visual, saliency, control network (CON) and default mode network (DMN). This suggests that the rare long-range exceptions are fundamental for the generation of the classic resting state networks. The boxplots of the figure use the standard Matlab convention, where the central mark indicates the median, while the bottom and top edges of the box indicate the 25th and 75th percentiles, respectively. The whiskers extend to the most extreme data points not considered outliers, which are plotted individually using a red ‘+’ symbol. We have added thee black stars, where the level of significance is higher than p<0.001 (Wilcoxon, rank sum).

**Figure 3B** shows boxplots of the errors of the FC fitting for the two models at the corresponding optimal working point. As hypothesised, the EDR+LR with the long-range connections perform significantly better in fitting the empirical data (p<0.001, Wilcoxon rank sum). This emphasises the important role of the long-range connections for brain function.

Importantly, as shown in the boxplots of **Figure 3C**, the two models are able to fit the level of turbulence in the empirical data (compare empirical, EDR+LR and EDR models). Turbulence is measured as the variability across time and space of the local synchronisation level (see Methods). In order to show the significance of turbulence in both empirical data and models, we constructed and computed turbulence for surrogate data shuffled while maintaining the spatiotemporal characteristics of the empirical data ^35^. The levels of turbulence of empirical data and models are significantly different (p<0.001, Wilcoxon rank sum) from surrogate data. This shows that EDR alone is enough to fit turbulence, which is not surprising given that other diffusive systems (i.e. with local coupling) such as fluids can also exhibit turbulence.

Having shown that the whole-brain model is able to fit the empirical data, **Figure 3D-H** shows the key experimental findings of how long-range exceptions enhance very specific aspects of information processing, namely information cascade, susceptibility, predictability and long-range connectivity (see Methods). Given that information transmission across space and time is mediated by the local level of synchronisation in brain vortex space, the first three measures are in this space. We also use a complementary measure of information processing given by the more conventional long-range functional correlation, that characterise the transmission of information over MNI space.

More specifically, our first measure of information cascade was motivated by the theory of turbulence, which recent research has found in human brain resting state dynamics ^28,29^. In particular, we investigated the *turbulent information cascade*, which measures the hierarchical transfer of information across scales. Briefly, turbulence can be defined by local levels of synchronisation at different spatial scales, *λ*, by means of the local order Kuramoto parameter *R*_*λ*_ (see Methods). This measure captures what we here call *brain vortex space, R*_*λ*,_ over time, inspired by the rotational vortices found in fluid dynamics, but of course not identical. The *information cascade flow* is the predictability of a given brain vortex space at scale *λ* from the brain vortex space at scale *λ* − Δ*λ* (where Δ*λ* is the discretisation of scale). In other words, the measure captures information transfer across scales through local synchronization in brain vortex space. Finally, we define the *information cascade* as the average of the *information cascade flow* across different scales (see Methods). In other words, enhancing the information cascade is a signature of enhanced information processing in terms of transferring information across spacetime. Note that this measure captures the transmission of information as mediated by the local level of synchronisation at different spatial scales.

As we hypothesised, the top panel of **Figure 3D** shows that the human EDR+LR model significantly increase the information cascade (p<0.001, Wilcoxon rank sum) compared to the EDR model. This is shown in more details in the bottom panel plotting the information cascade flow for the two models as a function of the spatial scale *λ*. Again, as hypothesised, we found an increase for the EDR+LR model (red line) compared with EDR models (blue line) as function of the scales in brain vortex space. These results demonstrate the functional importance of long-range connections for information processing.

Using whole-brain modelling also allowed us to test the importance of structural long-range exceptions for enhancing other information processing measures not in brain vortex space (and thus directly associated with turbulence). Specifically, we investigated the functional long-range connectivity (FC long-range), which is a measure of the average functional connectivity between pairs of brain regions over 40mm apart. **Figure 3E** shows a significant increase for FC long-range for the EDR+LR model (p<0.001, Wilcoxon rank sum). The increase in FC long-range for the EDR+LR model is concomitant with a smaller error in fitting to the full functional data (shown in **Figure 3B**).

Back in brain vortex space, yet another measure of information processing comes from measuring the susceptibility of the whole-brain model, which describes the reactivity and sensitivity of a system to external perturbations (see Methods). Systems with high susceptibility are more prone to encode external inputs and therefore important for cognition. As can be seen from **Figure 3F**, again the long-range connection model outperforms the simple EDR model. Thus, the long-range connections add sensitivity to external stimulation, potentially important for task-related information processing (see below).

Still in brain vortex space, we tested the predictability measure of information processing (see Methods) and found that the EDR+LR model outperforms the EDR model in terms of being better able to predict the future state of the brain (**Figure 3 G**).

Finally, we were interested in providing potential evidence for the involvement of the long-range exceptions in the generation of resting state networks. **Figure 3H** shows the level of fitting of both models with the seven Yeo resting state networks. As can be seen both models provide good fits to the resting state networks, but the EDR+LR model is significantly better at fitting the visual, saliency, control network (CON) and default mode network (DMN) (all p<0.001, Wilcoxon rank sum). This important result suggests that while the EDR could well be the underlying anatomical skeleton enabling resting state networks, the EDR+LR model enhances the generation of the resting state networks.

Overall, these findings rigorously confirm our hypothesis that rare long-range exceptions on top of the simple EDR wiring of the human brain significantly enhance information processing (as characterised by four independent measures).

### The role of turbulence for the ability of long-range exceptions to enhance information processing

Given that turbulence is present in the empirical brain dynamics and the two models, we investigated whether this could be facilitating the enhancement of information processing by rare long-range exceptions. In order to manipulate the level of turbulence, we turned to the simplest model system, namely that of a EDR ring architecture with and without LR.

Previous research has shown that creating a network with small-world properties such as the EDR+LR enhance the structural topological properties ^36^. Changing the topological brain anatomy has been studied in quite some details ^13-15^, while functional consequences of manipulating the underlying architecture have been studied less ^16,17^. However, the role of turbulence for information processing in architectures with rare long-range exceptions has not been studied.

As shown in **Figure 4A**, we constructed three different types of ring architectures: nearest neighbour (NN, black ring), EDR (blue ring) and rare long-range (EDR+LR, red ring). These formed the basis for three models of coupled Stuart-Landau oscillators, where we measured turbulence, averaged functional connectivity at long-range distances (FC long-range) and information cascade.

**Figure 4.**
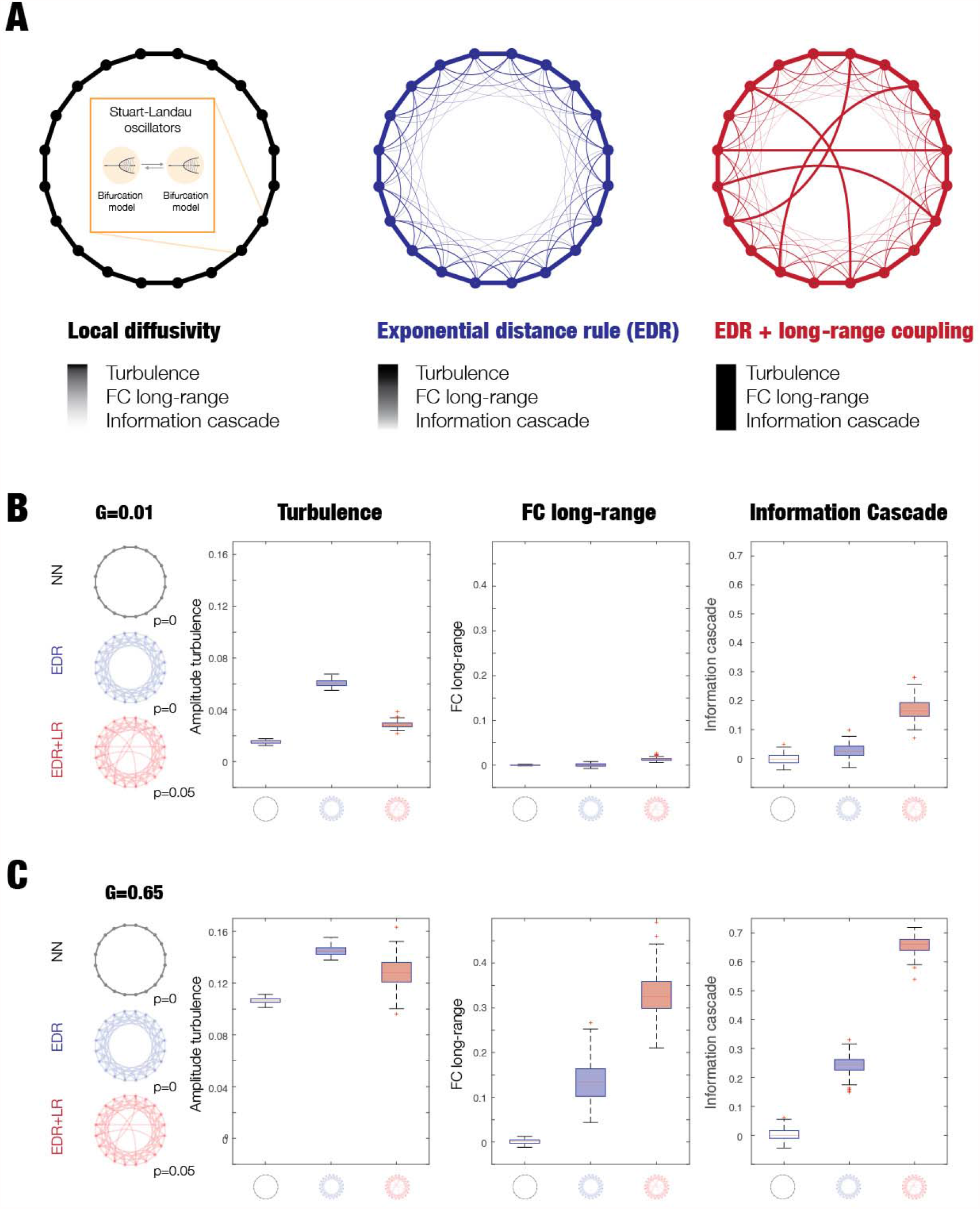
Using a simple ring structure to demonstrate the improvement in information processing by long-range exceptions is due to the underlying turbulence. **A)** We created three distinct ring architectures of coupling: nearest neighbour (NN, black ring), exponential distance rule (EDR, blue ring) and EDR with long-range exceptions (EDR+LR, red ring). In each architecture, we used Stuart-Landau oscillators to study the degree of turbulence, functional connectivity in long-range distances (FC long-range) and information cascade. **B)** We analysed the role of turbulence by varying the global coupling (G, see Results, Methods and Supplementary Figure 1) and how turbulence is amplifying the effects of the long-rare exceptions on FC long-range and information cascade. For a low-level turbulence with a low global coupling G=0.01, the figure shows three subpanels with a row of boxplots for the measures for the three ring architectures. As can be seen, at this low-level of turbulence regime, the effect of the long-range exceptions is only moderate. **D)** However, when we increase the level of turbulence of the EDR+LR model (G=0.65), the effect of the long-range exceptions is highly significant for both FC long-range (middle panel) and information cascade (rightmost panel). This shows clearly that turbulence combined with the specific EDR+LR architecture plays a significant role in enhancing the information cascade and information processing in general.

**Figure 4B** and **Figure 4C** show the role of turbulent fluctuations on the impact of long-range connections on information processing. **Figure 4B** shows the effect of low levels of turbulent fluctuations with a low global coupling G=0.01 in the three architectures. As can be seen from the the boxplots in the first panel, there are low levels of turbulent fluctuations for all three models, and comparatively modest effects of enhancing the effects of rare long-range exceptions on measures of FC long-range and information cascade (in the second and third panel).

In contrast, **Figure 4C** shows that increasing the global coupling to G=0.65 increase turbulent fluctuation levels in all models. Under these conditions, the effect of rare long-range exceptions is significantly amplified as shown by the measures for both FC long-range (middle panel) and information cascade (rightmost panel) (both p<0.001, Wilcoxon rank sum). These results confirm our hypothesis that turbulence provides the underlying functional regime allowing for the increase in information processing found in the EDR+LR model.

Furthermore, in **Supplementary Figure 1**, we investigated the dependence of these results as a function of global coupling and the probability of long-range exceptions in the model. **Figure S1A** shows these measures as a function of global coupling, G, in all three architectures. For the EDR+LR model, we used *p*=0.05 as the probability of long-range exceptions and found the largest effect was on FC long-range and information cascade (compare the non-overlapping red curves with the blue and grey curves for all values of G). This means that the level of long-range exceptions has a strong impact on information processing as revealed by the increase of functional correlations in the long-range connections and in the transmission of information reflected by the information cascade. It is interesting to note that in contrast to the other models, the NN model does not increase with G for the measures of FC long-range and information cascade. In contrast, the level of turbulence increases with G in all three architectures.

**Figure S1B** shows the effect of rewiring of long-range exceptions. For the optimal global coupling of the EDR+LR model, G=0.65, we plot the same three measures for all three architectures but now as a function of the probability of rewiring long-range exceptions. We observe the same effect as when systematically varying G and p=0.05 is close to optimal (see the red curve in the middle panel of **Figure S1B**).

Further insight on the low and high turbulent fluctuation regimes is provided by **Figure S1C**. Here, we investigated the EDR+LR model (with rewiring of long-range exceptions, p=0.05) in the low (G=0.01) and high (G=0.65) turbulent fluctuation regime. The first column shows snapshots of phases associated with the low (top) and high (bottom) turbulent fluctuation regime. As can be seen, phases are more clearly clustered in the high compared to low regime, reflecting local cluster synchronisation resembling turbulent vortices. The second column shows the distribution of the FC long-range for the low (top) and high turbulent fluctuation regimes (bottom); this reveals a strong increase of long-range FC across all pairs in the high compared to low regime.

Finally, we were able to show explicitly how information transmission across brain vortex space is influenced by long-range coupling. **Figure S1D** shows the *information cascade flow* (i.e. a measure *across* scales), rather than the information cascade (i.e. the *average* across scales) as shown in **Figure 3**. At optimal global coupling, there is a strong effect of long-range connectivity (compare red line, p=0.05 with blue line, p=0). The baseline (grey line) is added as a reference and corresponds to the information cascade of surrogates of the same timeseries where the time-ordering of phases were shuffled (100 repeats). We show the normalised information cascade flow (normalised with respect to baseline in the bottom plot), which shows exactly the same effect. Note in the normalised version, the flattening linear decay of information transfer for the model without long-range connections. In contrast, the EDR+LR model shows high information across all scales that decays slowly, explicitly reflecting the information cascade.

## Discussion

The brain appears to be unique in terms of its complex architecture spanning multiple scales ^37^. Unlike other known physical systems, where the elements communicate with nearest and close neighbouring elements (such as for example fluids or the heart), the brain uniquely possesses distant connections including a small contingent of long-range anatomical outliers, which – given their crucial role – we hypothesise have a significant role for enhancing information processing, presumably under strong evolutionary pressure. This hypothesis is difficult to test given that it is neigh impossible to isolate the long-range exceptions in animal experiments.

However, whole-brain modelling of empirical data offers a unique opportunity to test this hypothesis by creating models with and without rare long-range exceptions. A good definition from information processing in the brain is not universally agreed but here we provide four different complementary quantifications to characterise information processing from different perspectives. One simple way is to measure information processing by characterising the correlations between long-distance pairs of regions in MNI space; where a higher degree of correlation at long-range distance is indicative of transmission of information across space. A more sophisticated approach to fully characterise information transmission over space and time uses the concept of brain vortex space, defined as the local level of synchronisation and thus similar but not identical to rotational vortices found in fluid dynamics. Note that the key idea is that the level of local synchronisation (captured by the local Kuramoto parameter) is essential to the transmission of information across spacetime. Specifically, we used three different complementary measures measuring the information cascade, susceptibility and predictability.

These measures allowed us to confirm our hypothesis that the long-range exceptions confer a significant improvement in information processing using a large empirical neuroimaging dataset of 1003 human subjects. Specifically, we contrasted two whole-brain models, EDR and EDR+LR. In terms of function, the latter better fitted the functional empirical data and in addition showed a significant increase in information processing as measured by the specific measures of long-range FC and information cascade. Importantly, the EDR+LR model was also better able to fit the emergence of the classical resting-state networks, providing potentially a link to cognition.

Our quantification of information processing were inspired by the pioneering work of Yoshiki Kuramoto, who demonstrated turbulence using coupled oscillators to describe the rich variability of local synchronisation ^30^. As such these measures do not depend on turbulence per se to measure information processing. We were, however, interested to see whether a turbulent regime enhance information processing through the rare long-range exceptions.

To this end, we studied models using the simplest possible ring architectures. We were able to demonstrate that all three models with different ring architectures of coupled oscillators (with nearest neighbour, EDR and EDR+LR) are capable of supporting a turbulent fluctuation regime but are not all equally efficient for information transfer. By varying the level of turbulence, we were able to show that high levels of turbulence the EDR+LR model significantly amplifies information processing as reflected in a concomitant significant increase of long-range FC, and information cascade (defined as a measure of information flow providing predictability across scales). These results demonstrate the potentially immense functional benefits for information processing in the turbulent fluctuation regime sustained by a brain architecture with rare long-range exceptions.

Beyond the demonstration that rare long-range exceptions amplify information processing due to the underlying turbulence, we note two important complementary facts. First, as shown by Andrey Kolmogorov, turbulence in fluid dynamics is providing the basis for highly efficient energy transmission. On an abstract level, information transfer is analogous to energy transfer, and in fact mathematical research has shown the close links between propagation of disturbances and the transmission of information ^38,39^. Kolmogorov also showed that turbulence is characterised by power laws revealing a cascade of energy and information ^40,41^ (see excellent review in ^42^).

Second, we have recently shown that human brain dynamics are turbulent in the sense that similar power laws are found Kolmogorov’s ‘structure-functions’, which corresponds to the functional connectivity as a function of the distance. In other words, the existence of a power law for human brain dynamics is not only evidence of turbulence but also evidence for the efficient transmission of correlations, i.e. transmission of information across spacetime. Strengthening this line of evidence, we also used the complementary Kuramoto framework, showing rich variability of the local level of synchronisation across spacetime in the empirical brain imaging data. Equally, fitting the empirical data with a whole-brain Hopf model based on Stuart-Landau oscillators, we were able to causally show that at the optimal working point, turbulence is maximal and associated with efficient information transmission ^28^.

Beyond the fundamental properties of information processing, it is of interest to note that the theoretical results from the model with the ring architecture shows that small proportion of long-range connections improves the information cascade (independent of their spatial location). However, the empirical results using a model with the brain architecture clearly show that these long-range exceptions are not spatially random but closely linked to the emergence of functionally important resting-state networks. Given that these networks have been shown to play a key role in task-based processing ^43-45^, we speculate that this suggests that evolution has improved on the basic EDR by refining long-range exceptions thereby improving brain function most notably its ability to perform certain behaviours thereby optimising survival. This will need to be further explored in cross-species investigations.

Research has investigated the hierarchical organisation of brain structure and function ^7,46-48^, which has demonstrated a common network available for the orchestration of task and rest ^28^. Previous research has demonstrated that the turbulent core of brain activity is largely overlapping with brain regions known to be involved in lower level sensory processing ^28^. We therefore hypothesised that the anatomical basis of this orchestration could be well served by the long-range outliers, which effectively are able to control brain activity in the turbulent core sustained by the underlying EDR. An obvious next step would therefore be to study a task in a large empirical dataset, a challenging prospect given the difficulty at the required granularity for whole-brain models to capture the dynamical complexity specific to a given task.

Structural analysis of the cortical network has emphasised its spatial embedding, showing that weight and distance of connections are tightly interwined. Connection weights span over five orders of magnitude and the action of the EDR means that the average connection weight declines exponentially with distance ^11^. A corollary of the decline in weight is that neighbouring areas show 100% connectivity with connection densities falling to very low levels with distance. These considerations suggest that long-distance connections confer a high degree of binary specificity to the cortical network which is amply confirmed by statistical analysis ^1^. Further, this analysis shows that long-distance connections carry precise signatures that ensure an important role of globalisation to a small group of areas. An intriguing finding of the present study is that the weight values of long-range connections appear to play a decisive role, given the marked differences between EDR and EDR+LR models in supporting the turbulent-like dynamics of information processing.

The present study provides a potential framework for explaining the relatively fast speed of computations needed for survival of the individual and species, which requires the fast interaction between feedforward and feedback brain connections. Given that the extraordinary slow average transmission delay between neurons, typically on the order of 40 ms between neurons, it has long been a conundrum how the brain can quickly distinguish between different categories of stimuli ^49^. Take for example the neuroimaging studies using magnetoencephalography (MEG) which have shown activity around 130-170 ms in the fusiform face area (FFA) when faces are presented ^50^. This rapid processing is likely potentiated by scale-free network processing in the turbulent core in the sensory regions and could potentially be a purely feedforward phenomenon. Crucially, however, human neuroimaging experiments have shown that the feedback provided by the long-range exceptions must play a key role in directing the flow of information ^51,52^. As an example, MEG studies of infant and adult faces have shown activity in the FFA for both stimuli but with simultaneous activity in the orbitofrontal cortex (OFC) at around 130ms only for the infant faces ^52^. Interestingly, even small deviations from the infant face template such as cleft lip, leads to much diminished activity in the OFC ^53^. This ‘parental instinct’ is found in even non-parents and clearly plays a role in directing attention to the special category of infants, presumably to ensure that we provide the necessary caregiving, even when we are not the parents ^54^. The long-range feedback from the OFC, presumably via the inferior longitudinal fasciculus, is controlling the rapid information processing flow, prioritising infant faces ^52,53^ and sounds ^55^ over other less important stimuli.

The study here relied on human brain neuroimaging which is a relatively coarse and slow method. In other words, the space and time scales analysed here are limited to the order of mm and seconds, respectively. Complementary to this approach, it would be of considerable interest to study the Bauplan of the brain at a cellular level or circuit level. It is well-known that cell-to-cell (rather than region-to-region or even voxel-to-voxel) interactions on a very fine spatial scale can influence the dynamics of the brain even on relatively long the time scales (seconds) relevant for the dynamical measures considered here ^56,57^. Progress has been made at finer scales as shown by the discovery of the signature of turbulence at the circuit level in the rodent hippocampus ^29^.

The crucial functional role played by the rare long-range exceptions under a turbulent regime has important implications for the hierarchical organisation of brain processing. One function of the EDR is that it confers a core-periphery structure in the cortex, where the core is largely centred on the prefrontal cortex ^2,11^. The high-efficiency cortical core has been speculated to provide the structural underpinnings of the Global Workspace hypothesis ^58,59^, which proposes that recurrent processing in the core allows amplification and globalization of conscious states ^2,47,60^. These results also provide important underpinnings for other theories of consciousness such as the Integrated Information Theory (IIT) ^61^ and the Temporo-spatial Theory of Consciousness (TTC) ^62^, given that the turbulent-like regime promotes the efficient information cascade needed for spatiotemporal integration. A better and more detailed investigation of the role of particular EDR outliers is expected to give an improved understanding of the link between structure and higher cognitive function. Furthermore, since the cortical core as defined by the EDR is found across species ^10-12,63^, the exploration of its involvement in turbulence across species would lead to a better understanding of comparative cognitive function.

The shaping of functional activity by a fixed anatomical structure is a conundrum that brings to mind Thomas Aquinas famous dictum: “*Quidquid recipitur ad modum recipientis recipitur*”, i.e. container (or recipient) shapes the content. It has been proposed that the flexibility of brain function associated with the rich palette of behaviours is linked to changeable connectivity through for example neuromodulation ^64,65^. This *effective connectivity* is known also to change with brain states during wakefulness, as well as in light and deep sleep. Thus, it would of considerable interest to investigate how the information cascade changes in different brain states.

In addition, we hypothesise that turbulence, information cascade and especially the lack of control of these may play a central role in neuropsychiatric disorders. The present framework would lend itself well to causally describe the emotional and information processing changes found in neuropsychiatric disorders and may provide a novel way to find sensitive and specific biomarkers.

These findings also pose a question regarding the role of rare long-range exceptions across species. Future research could investigate the causal link between the underlying brain architecture of different species, dynamical working point, turbulence, information cascade and behavioural complexity. Ultimately, this could help cast new light on the deep question what makes us human.

## STAR Methods

### RESOURCE AVAILABILITY

#### Lead Contact

Further information and requests for resources should be directed to and will be fulfilled by the Lead Contact: Morten L. Kringelbach.

#### Materials Availability

The data set used for this investigation was from an independent publicly available dataset of fMRI data, where we chose a sample of 1003 participants selected from the March 2017 public data release from the Human Connectome Project (HCP).

#### Data and Code Availability

The HCP dataset is available at https://www.humanconnectome.org/study/hcp-young-adult. The code to run the analysis is available on GitHub (https://github.com/decolab/cb-longrange).

#### Experimental models and subject details

##### Neuroimaging Ethics

The Washington University–University of Minnesota (WU-Minn HCP) Consortium obtained full informed consent from all participants, and research procedures and ethical guidelines were followed in accordance with Washington University institutional review board approval.

##### Neuroimaging Participants

The data set used for this investigation was selected from the March 2017 public data release from the Human Connectome Project (HCP) where we chose a sample of 1003 participants.

### METHOD DETAILS

#### Neuroimaging acquisition for fMRI HCP

The 1003 HCP participants were scanned on a 3-T connectome-Skyra scanner (Siemens). We used one resting state fMRI acquisition of approximately 15 minutes acquired on the same day, with eyes open with relaxed fixation on a projected bright cross-hair on a dark background. The HCP website (http://www.humanconnectome.org/) provides the full details of participants, the acquisition protocol and preprocessing of the data for resting state.

#### Preprocessing and extraction of functional timeseries in fMRI resting data

The preprocessing of the HCP resting state and task datasets is described in details on the HCP website. Briefly, the data is preprocessed using the HCP pipeline which is using standardized methods using FSL (FMRIB Software Library), FreeSurfer, and the Connectome Workbench software ^66,67^. This preprocessing included correction for spatial and gradient distortions and head motion, intensity normalization and bias field removal, registration to the T1 weighted structural image, transformation to the 2mm Montreal Neurological Institute (MNI) space, and using the FIX artefact removal procedure ^67,68^. The head motion parameters were regressed out and structured artefacts were removed by ICA+FIX processing (Independent Component Analysis followed by FMRIB’s ICA-based X-noiseifier ^69,70^). Preprocessed timeseries of all grayordinates are in HCP CIFTI grayordinates standard space and available in the surface-based CIFTI file for each participants for resting state.

We used a custom-made Matlab script using the ft_read_cifti function (Fieldtrip toolbox ^71^) to extract the average timeseries of all the grayordinates in each region of the Schaefer parcellation, which are defined in the HCP CIFTI grayordinates standard space. Furthermore, the BOLD time series were transformed to phase space by filtering the signals in the range between 0.008-0.08 Hz, where we chose the typical highpass cutoff to filter low-frequency signal drifts ^72^, and the lowpass cutoff to filter the physiological noise, which tends to dominate the higher frequencies ^72,73^. We then applied the Hilbert transforms in order to obtain the phases of the signal for each brain node as a function of the time. We computed the functional connectivity (FC) as the correlation between the BOLD timeseries in all 1000 regions in the Schaefer Parcellation.

#### Structural connectivity using dMRI

The Human Connectome Project (HCP) database contains diffusion spectrum and T2-weighted imaging data from 32 participants with the acquisition parameters described in details on the HCP website ^74^. The freely available Lead-DBS software package (http://www.lead-dbs.org/) provides the preprocessing which is described in details in Horn and colleagues ^75^ but briefly, the data was processed using a generalized q-sampling imaging algorithm implemented in DSI studio (http://dsi-studio.labsolver.org). Segmentation of the T2-weighted anatomical images produced a white-matter mask and co-registering the images to the b0 image of the diffusion data using SPM12. In each HCP participant, 200,000 fibres were sampled within the white-matter mask. Fibres were transformed into MNI space using Lead-DBS ^76^. We used the standardized methods in Lead-DBS to produce the structural connectomes for both Schaefer 1000 parcellation Scheme ^77^ where the connectivity has been normalised to a maximum of 0.2.

#### Exponential Distance Rule

We fitted the underlying anatomy obtained with human diffusion MRI (see above) to the *Exponential Distance Rule (EDR)*, originally derived from massive retrograde tract tracing in non-human primates ^11^. Mathematically this can be expressed as an exponential decay function:

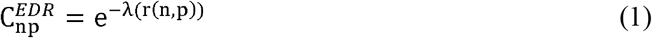

where r(n, p) is the Euclidean distance between the regions n and p, and the decay, *λ*.

#### Schaefer parcellation

Schaefer and colleagues created a publicly available population atlas of cerebral cortical parcellation based on estimation from a large data set (N = 1489) ^77^. They provide parcellations of 400, 600, 800, and 1000 areas available in surface spaces, as well as MNI152 volumetric space. We used here the Schaefer parcellation with 1000 areas and estimated the Euclidean distances from the MNI152 volumetric space and extracted the timeseries from HCP using the HCP surface space version.

### QUANTIFICATION AND STATISTICAL ANALYSIS

#### Hopf Whole-Brain Model

At the heart of whole-brain network models is the link between anatomical structure and functional dynamics, introduced more than a decade ago ^78,79^. Typically, the anatomy is represented by the structural connectivity (SC) of an individual or average brain, measured *in vivo* by diffusion MRI (dMRI) combined with probabilistic tractography. The spatial resolution is in the order of 1-2 mm, but with ultra-high field MRI resolutions 0.4 mm can be reached. The structural connectome denotes the wire-diagram of the connections between cortical regions as ascertained from dMRI tractography. The functional global dynamics result from the mutual interactions of local node dynamics coupled through the underlying empirical anatomical SC matrix. Whole-brain models aim to balance between complexity and realism in order to describe the most important features of the brain *in vivo* ^18^. Recent developments have shown that whole-brain models are able to describe not only static FC (averaged over all time points), but also dynamical measurements like the temporal structure of the activity fluctuations, the so-called functional connectivity dynamics (FCD) ^24,80^.

Here we used the Hopf whole-brain model consisting of coupled dynamical units (ROIs or nodes) representing the N cortical brain areas from a given parcellation ^24^. We used all 1000 cortical nodes in the Schaefer parcellation. The local dynamics of each brain region is described by the normal form of a supercritical Hopf bifurcation, also known as the Landau-Stuart Oscillator, which is the canonical model for studying the transition from noisy to oscillatory dynamics ^81^. Coupled together with the brain network architecture, the complex interactions between Hopf oscillators have been shown to reproduce significant features of brain dynamics observed in electrophysiology^82,83^, MEG ^84^ and fMRI ^85,86^.

The whole-brain dynamics was defined by the following set of coupled equations in Cartesian coordinates:

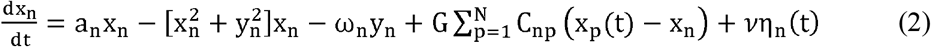

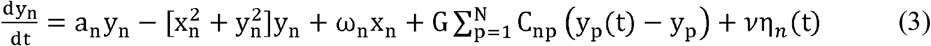

where η_n_(t) is additive Gaussian noise with standard deviation *υ = 0*.*01*. This normal form has a supercritical bifurcation a_n_ =0, so that if a_n_ >0, the system engages in a stable limit cycle with frequency *f_n_* = ω_n_ /2*π*. On the other hand, when a_n_ <0, the local dynamics are in a stable fixed point representing a low activity noisy state. Within this model, the intrinsic frequency *f*_*n*_ is estimated from the empirical data as the peak of the power spectrum. Here, the subindex n denotes the region taken from (1..N), where N is the total number regions. The local bifurcation parameters, a_n =_ -0.02, are at the brink of the local bifurcations where the best fitting to the empirical data was demonstrated. The variable x_n_ emulates the BOLD signal of each region n. To model the whole-brain dynamics we added an additive coupling term representing the input received in region n from every other region p, which is weighted by the corresponding structural connectivity C_np_, which can correspond to the connectivity with EDR 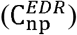 or 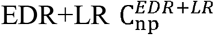. In this term, G denotes the global coupling weight, scaling equally the total input received in each brain area. All the measures related to whole-brain model were estimated for each global coupling work point, G, running the simulations 100 times and averaging the results.

#### Turbulence

The study of turbulence in modern physics has led to enormous progress in the fields of fluid and oscillator dynamics ^30,38,42^. It is well-known that coupled oscillators fit brain dynamics ^24,82,83^. However, for the brain to function optimally, it has to support efficient information transfer and here we hypothesise that turbulence provides the intrinsic backbone necessary for brain dynamics.

Historically, in fluid dynamics the study of turbulence was greatly influenced by Richardson’s concept of cascading eddies reflecting energy transfer, where the hierarchical organisation of different sizes of eddies is schematised for the turbulent so-called ‘inertial subrange’, i.e. the range where turbulence kinetic energy is transferred without loss from large to smaller scales. Here, we propose that this hierarchical organisation of oscillators leads to an information cascade in the inertial subrange.

The measure of **turbulence** comes from the study of coupled oscillators and in particular the seminal studies by Kuramoto investigating turbulence in coupled oscillators ^30^. Specifically, in a coupled oscillator framework, the Kuramoto local order parameter represents a spatial average of the complex phase factor of weighted coupling of local oscillators.

Specifically, the amplitude turbulence, 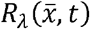, is defined as the modulus of the local order parameter for a given brain node as a function of time:

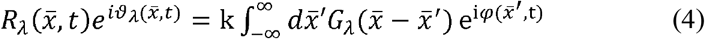

where 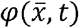 are the phases of the spatiotemporal data, *G*_*λ*_ is the local weighting kernel 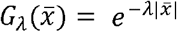, k is the normalisation factor 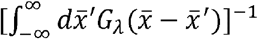 and *λ* is defines the scaling of local weighting. In other words,*R*_*λ*_ defines local levels of synchronisation at a given scale, *λ*, as function of space, 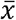, and time, *t*, This measure captures what we here call *brain vortex space, R*_*λ*_, over time, inspired by the rotational vortices found in fluid dynamics, but of course not identical.

The level of amplitude turbulence is defined as the standard deviation of the modulus of Kuramoto local order parameter and can be applied to the empirical data of any physical system. The amplitude turbulence, *D*, corresponds to the standard deviation across time and space of *R*_*λ*_:

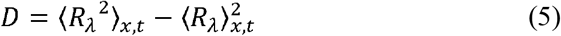

where the brackets ⟨ ⟩_*x,t*_ denotes averages across space and time.

Except for a very recent study ^28^, amplitude turbulence has been studied exclusively in the supercritical regime of the Stuart-Landau coupled oscillators ^87,88^. Yet, given that a range of papers have demonstrated that the Stuart-Landau oscillators fit the neuroimaging brain data when used in the subcritical regime at the edge of the bifurcation ^24,84,85,89^, it was recently demonstrated to fit the functional MRI data from 1003 participants and to exhibit amplitude turbulence as characterised by Equation 5 ^28^. The subcritical regime at the edge of the bifurcation is not showing deterministic spatiotemporal chaos in the way found in under the supercritical regime but rather some very interesting state dynamics, perhaps best described as turbulent-like or turbulent fluctuation, which are described by Equation 5.

#### Measures of Information Processing

Here we quantify information processing by characterising the transmission of information over spacetime. First, we defined a conventional method, namely 1) long-range functional connectivity, i.e. correlations in MNI space but only taking into account long range Euclidean distances. Secondly, we defined three complementary information processing transmission measurements in brain vortex space: 2) information cascade, 3) susceptibility and 4) predictability. The concept of brain vortex space is given by the local level of synchronisation and the transmission of information over space and time. This is important given that information is transferred across space and time, mediated by local synchronisation.

#### Long-range functional connectivity

Rather than working in brain vortex space, it is also desirable to characterize information processing by using conventional functional connectivity as a function of a given distance, given by FC(r) in MNI space. In fact, this can be derived from Kolmogorov’s concept of *structure function* for a given distance, *S(r)*, of a variable *u* ^40,41^ in spatiotemporal data in the following manner:

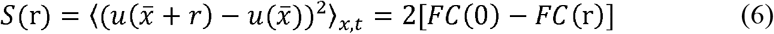

In this equation the basic functional connectivity, *FC*, between two points separated by the Euclidean distance *r*, is given by:

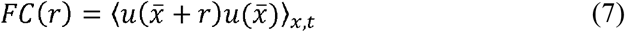

where the symbol ⟨ ⟩_*x,t*_ refers to the average across the spatial location 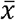 of the nodes and time. Using these relations, we are able to study the impact of different architectures on the functional connectivity for long-range distances (FC long-range), here defined as distances over 40mm.

#### Information cascade

As mentioned, the measure of information cascade uses the concept of *brain vortex space, R_λ_*, and is defined as the average across scales, *λ*, of the *information cascade flow*, which indicates the predictability of a given brain vortex space at scale *λ* from the brain vortex space at scale *λ* − Δ*λ* (where Δ*λ* is the discretisation of scale).

In other words, the measure captures information transfer across scales through local synchronization in brain vortex space. Mathematically, *information cascade flow* can be expressed as:

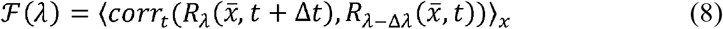

where Δ*t*, is the size of the time step (and thus fixed), and Δ*λ* is also fixed according to the used range

*λ* = [4 2 1 0.5 0.25 0.125 0.0625 0.0312]. The symbol ⟨ ⟩_*x*,_ refers to the spatial average of the correlation over time, *corr*_*t*_ (*R*_*λ*_, *R*_*λ* − Δ *λ*_). The correlation of the level of synchronisation at different scales and consecutive time captures the flow of transmission of information across spacetime. In particular we wanted to compress this into the single measure *information cascade, I*, which is simply the average of . ℱ(*λ*) across different scales (*λ*):

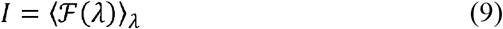

where the symbol ⟨ ⟩_*λ*_ refers to the average across *λ*.

#### Susceptibility

Another way of characterising information processing is to measure the sensitivity of the system when it is perturbed. This requires building a whole-brain model of the system. The advantage of using a model means that we can easily measure the sensitivity of the model to external perturbation, which is a standard physics measure usually called *susceptibility*. Indeed, this measure can be obtained by perturbing the whole-brain model at the optimal working point by randomly changing the local bifurcation parameter, a_n_, in the range [-0.02:0] at the node level n. Susceptibility is measured by the effects of the perturbation on the brain vortex space, i.e. estimated by measuring the modulus of the local Kuramoto order parameter, i.e. 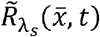 for the perturbed case, and 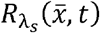 for the unperturbed case. Thus, we define susceptibility s follows:

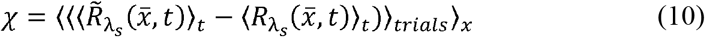

where ⟨ ⟩_*t*,_ ⟨ ⟩_*trails*_ and ⟨ ⟩_*x*_ are the mean averages across time, trials and space, respectively.

#### Predictability in brain vortex space

Yet another measure of information processing is to measure the level of predictability in brain vortex space for *q* steps in the future. The predictability is computed by the correlation 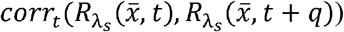, where *corr*_*t*_ signifies correlation across time.

#### Ring architecture

In order to understand the consequences of long-range exceptions on dynamics in the context of turbulence, we study the simplest system, namely a ring architecture. We constructed three different types of ring architectures: nearest neighbour (NN, black ring), EDR (blue ring) and rare long-range (LR, red ring) in the following way:

1. Nearest neighbour (NN) ring architecture, and where 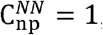, if p=n±1 or 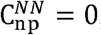, otherwise.
2. The EDR ring architecture is given by Equation 1, here called 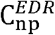. Here r(n, p) is the distance between nodes n and p, given by the absolute value divided by a scaling factor *k*=10, i.e. |p − n|*/k*. The coupling decay factor is given by λ = 1.
3. Rare long-range (LR) is similar to the EDR ring architecture but with additional random connections, here called 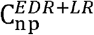. We add these connections similar to the small-world procedure described in Watts and Strogatz ^36^ by going over all connections and replacing pairs randomly with a probability of *p* and allocating these connections a coupling strength of 0.25.

This allowed us to create three corresponding models of coupled Stuart-Landau oscillators, where for various scenarios we measured turbulence, averaged functional connectivity at long-range distances (FC long-range) and information cascade. We ran the three models for 1000 timesteps over 100 trials. Specifically, turbulence was computed as above, while the FC long-range was computed as the mean value of functional connectivity pairs with the top 20% largest distance. The range of scales of *λ* used for estimating . ℱ and *I* was given by λ = [4 2 1 0.5 0.25 0.125 0.0625 0.0312] (as above).

## Author contributions

G.D. and M.L.K. designed the study, developed the method, performed analyses, wrote and edited the paper.

## Acknowledgements

We would like to acknowledge the important theoretical contributions made by Profs Yoshiki Kuramoto, Hiroya Nakao, Ryan Raut and Marcus Raichle. G.D. is supported Spanish national research project (ref. PID2019-105772GB-I00 MCIU AEI) funded by the Spanish Ministry of Science, Innovation and Universities (MCIU), State Research Agency (AEI); HBP SGA3 Human Brain Project Specific Grant Agreement 3 (grant agreement no. 945539), funded by the EU H2020 FET Flagship programme; SGR Research Support Group support (ref. 2017 SGR 1545), funded by the Catalan Agency for Management of University and Research Grants (AGAUR); Neurotwin Digital twins for model-driven non-invasive electrical brain stimulation (grant agreement ID: 101017716) funded by the EU H2020 FET Proactive programme; euSNN European School of Network Neuroscience (grant agreement ID: 860563) funded by the EU H2020 MSCA-ITN Innovative Training Networks; CECH The Emerging Human Brain Cluster (Id. 001-P-001682) within the framework of the European Research Development Fund Operational Program of Catalonia 2014-2020; Brain-Connects: Brain Connectivity during Stroke Recovery and Rehabilitation (id. 201725.33) funded by the Fundacio La Marato TV3; Corticity, FLAG–ERA JTC 2017 (ref. PCI2018-092891) funded by the Spanish Ministry of Science, Innovation and Universities (MCIU), State Research Agency (AEI). MLK is supported by the Center for Music in the Brain, funded by the Danish National Research Foundation (DNRF117), and Centre for Eudaimonia and Human Flourishing funded by the Pettit and Carlsberg Foundations.

## Figures

**Supplementary Figure 1.**
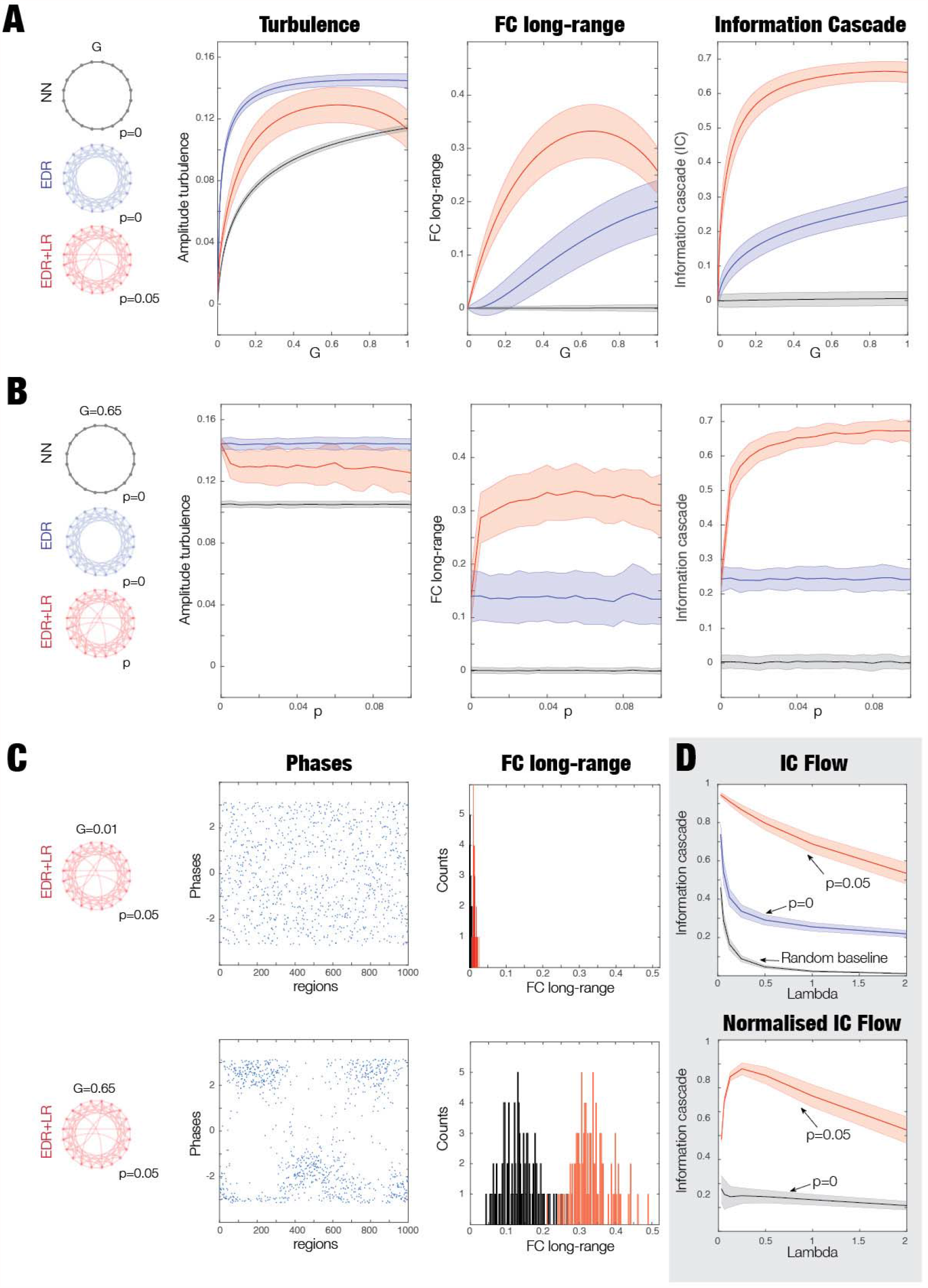
Comparing the measures of turbulence, FC long-range and information cascade in different ring architectures. **A)** First, we plot these measures as a function of global coupling, G, in the models. In the EDR+LR model, p=0.05, as the probability of long-range connections. The EDR+LR model is the best, showing long-range connectivity having a large effect on FC long-range and information cascade (compare the red curves with the blue and grey curves). **B)** Using the optimal global coupling of the EDR+LR model, G=0.65, we plot the same three measures but now as a function of the probability of long-range connections. Again, we find that the EDR+LR model is the best (compare the red curves with the blue and grey curves). **C)** We were interested in comparing low and high turbulence regime and investigated the EDR+LR model with G=0.01 (low turbulence), and G=0.65 (high turbulence regime) in both cases in p=0.05. The figure shows a column with snapshots of phases in the low turbulent regime (top) and high turbulent regime (bottom). The phases are clearly more clustered for the high compared to low turbulent regime. The second column shows the distribution of the FC long-range for the low turbulent regime (top) and high turbulent regime (bottom). Again, note the strong increase of the FC long-range for the high compared to low turbulent regime. **D)** We show the information cascade flow (IC Flow) for the three different ring architectures at the optimal global coupling, revealing the effect of the long-range connectivity. The bottom is a normalised version of the top plot with respect to the baseline (see text). This demonstrates explicitly how the information transmission across the scale of the brain vortex space is affected by the long-range coupling.

